# Multi-Parametric and Multi-Regional Histogram Analysis of MRI: Revealing Imaging Phenotypes of Glioblastoma Correlated with Patient Survival

**DOI:** 10.1101/235861

**Authors:** Chao Li, Shuo Wang, Angela Serra, Turid Torheim, Jiun-Lin Yan, Natalie R Boonzaier, Tomasz Matys, Mary A McLean, Florian Markowetz, Stephen J Price

## Abstract

**Introduction:** Glioblastoma is characterized by its remarkable heterogeneity and dismal prognosis. Histogram analysis of quantitative magnetic resonance imaging (MRI) is an important *in vivo* method to study intratumoral heterogeneity. With large amounts of histogram features generated, integrating these modalities effectively for clinical decision remains a challenge.

**Methods:** A total of 80 patients with supratentorial primary glioblastoma were recruited. All patients received surgery and standard regimen of temozolomide chemoradiotherapy. Diagnosis was confirmed by pathology. Anatomical T2-weighted, T1-weighted post-contrast and FLAIR images, as well as dynamic susceptibility contrast (DSC), diffusion tensor imaging (DTI) and chemical shift imaging were acquired preoperatively using a 3T MRI scanner. DTI-p, DTI-q, relative cerebral blood volume (rCBV), mean transit time (MTT) and relative cerebral blood flow (rCBF) maps were generated. Contrast-enhancing (CE) and non-enhancing (NE) regions of interest were manually delineated. Voxel intensity histograms were constructed from the CE and NE regions independently. Patient clustering was performed by the Multi-View Biological Data Analysis (MVDA) approach. Kaplan-Meier and Cox proportional hazards regression analyses were performed to evaluate the relevance of the patient clustering to survival. The histogram features selected from MVDA approach were evaluated using receiver operator characteristics (ROC) curve analysis. The metabolic signatures of the patient clusters were analyzed by multivoxel MR spectroscopy (MRS).

**Results:** The MVDA approach yielded two final patient clusters, consisting of 53 and 27 patients respectively. The two patient subgroups showed significance for overall survival (p = 0.007, HR = 0.32) and progression-free survival (p < 0.001, HR = 0.33) in multivariate Cox regression analysis. Among the features selected by MVDA, higher mean value of DTI-q in the non-enhancing region contributed to a worse OS (HR = 1.40, p = 0.020) and worse PFS (HR = 1.36, p = 0.031). Multivoxel MRS showed N-acetylaspartate/creatine (NAA/Cr) ratio between the two clusters, both in the CE region (p < 0.001) and NE region (p = 0.013). Glutamate/Cr (Glu/Cr) ratio and glutamate + glutamine/Cr (Glx/Cr) of the cluster 1 was significantly lower than cluster 2 (p = 0.037, and 0.027 respectively) In the NE region.

**Discussion:** This study demonstrated that integrating multi-parametric and multi-regional MRI histogram features may help to stratify patients. The histogram features selected from the proposed approach may be used as potential imaging markers in personalized treatment strategy and response determination.

## Introduction

Glioblastoma represents the most common primary brain malignant tumours in adults, characterized by its dismal prognosis [1]. The remarkable heterogeneity of glioblastoma may cause the inconsistent treatment response of patients. There is a rising need for validated markers to assess interpatient variability, plan personalized treatment and predict treatment response. With recent advances in molecular biology, the diagnostic and/or prognostic significance of genetic markers is established [2, 3]. However, the assessment of these genetic markers relies on invasive biopsies or resections and may be prone to sampling errors.

Magnetic resonance imaging (MRI) shows potential in capturing imaging features non-invasively prior to treatment. The features extracted from MRI was shown to be able to reveal phenotypes with different molecular pathways and survivals [4]. Histogram features extracted from the quantitative MRIs can characterize tumor heterogeneity by measuring spatial variation within the whole tumor, which may be related with tumor malignancy and patient survival [5]. Due to the emergence of multiple MRI sequences, considerable amounts of histogram features can be generated from the multiple MRI modalities. However, integrating these modalities effectively and selecting optimal histogram features for clinical decision remains a challenge.

Although machine learning algorithms have achieved success in selecting relevant image features and stratifying patients [6], classical machine learning techniques that work on a single imaging modality may be ineffective on multiple imaging modalities. Therefore, new machine learning methodologies for patient stratification were needed. Given the success of multi-view learning techniques in genomic studies, the purpose of this current study was to explore the multi-parametric and multi-regional histogram features with a previously reported multi-view feature selection and clustering methods [7]. This multi-view method was developed to jointly analyze multiple genomic features. We hypothesized that it can be applied to multiple imaging modalities for more stable clustering results and better insights into the patient characterization [8, 9]. The patient subgroups identified by this method may display different outcomes and different metabolic signatures.

The advanced MRI sequences we used include perfusion and diffusion imaging, which may confer physiological information and compensate the non-specificity of conventional imaging. Dynamic susceptibility contrast (DSC) MRI is one of the most commonly-used perfusion techniques, which measures the kinetics of contrast agent passing through the capillary bed [10]. Several biomarkers, including the relative cerebral blood volume (rCBV), mean transit time (MTT) and relative cerebral blood flow (rCBF) are estimated from the kinetics curve. Diffusion tensor imaging (DTI) is a method which may detect the tumor infiltration by measuring the magnitude and direction of water molecule movement [11]. To interpret the high-dimensional tensor imaging, a decomposition into isotropic component (p) and anisotropic component (q) was proposed [12]. This method measures the isotropic and anisotropic diffusion of water molecules and has shown its utility in predicting tumor progression [13] and patient survival [14]. More recently, integrating these modalities was considered useful in revealing the intratumoral invasive component [15] and predicting IDH genotype in high-grade gliomas [16]. In this study, we used the histogram features extracted from above multi-parametric and multi-regional quantitative MRI to stratify patient groups and assess the relevance to treatment outcome.

## Methods

### Patients

From July 2010 to August 2015, patients with supratentorial primary glioblastoma were prospectively recruited. Patients who had a history of previous brain tumor, cranial surgery, radiotherapy/chemotherapy, or contraindication for MRI scanning were excluded. This study was approved by the local institutional review board. Signed informed consent was obtained from each patient.

A total of 80 patients were included into the study. All patients had good performance status (World Health Organization performance status 0-1) before surgery. Neuronavigation (StealthStation, Medtronic) and 5-aminolevulinic acid fluorescence were used to guide surgery for maximal safe resection. Standard regimen of temozolomide chemoradiotherapy was performed after surgery when patients were stable. Extent of resection was assessed according to the postoperative MRI scans within 72 hours, classified as gross total resection, subtotal resection or biopsy of the contrast enhancement. Patients’ treatment response was evaluated according to the Response Assessment in Neuro-oncology criteria [17].

### MRI Acquisition

All MRI sequences were performed at a 3-Tesla MRI system (Magnetron Trio; Siemens Healthcare, Erlangen, Germany) with a standard 12-channel receive-head coil. MRI sequences were acquired as following: post-contrast T1-weighted sequence (TR/TE/TI 2300/2.98/900 ms; flip angle 9°; FOV 256 × 240 mm; 176-208 slices; no slice gap; voxel size 1.0 × 1.0 × 1.0 mm) after intravenous injection of 9 mL gadobutrol (Gadovist,1.0 mmol/mL; Bayer, Leverkusen, Germany); T2-weighted sequence (TR/TE 4840-5470/114 ms; refocusing pulse flip angle 150°; FOV 220 × 165 mm; 23-26 slices; 0.5 mm slice gap; voxel size of 0.7 × 0.7 × 5.0 mm); T2-weighted fluid attenuated inversion recovery (FLAIR) (TR/TE/TI 7840-8420/95/2500 ms; refocusing pulse flip angle 150°; FOV 250 × 200 mm; 27 slices; 1 mm slice gap; voxel size of 0.78125 × 0.78125 × 4.0 mm). PWI was acquired with a dynamic susceptibility contrast-enhancement (DSC) sequence (TR/TE 1500/30 ms; flip angle 90°; FOV 192 × 192 mm; FOV 192 × 192 mm; 19 slices; slice gap 1.5 mm; voxel size of 2.0 × 2.0 × 5.0 mm;) with 9 mL gadobutrol (Gadovist 1.0 mmol/mL) followed by a 20 mL saline flush administered via a power injector at 5 mL/s. DTI was acquired with a single-shot echoplanar sequence (TR/TE 8300/98 ms; flip angle 90°; FOV 192 × 192 mm; 63 slices; no slice gap; voxel size 2.0 × 2.0 × 2.0 mm). Multivoxel 2D ^1^H-MRS chemical shift imaging (CSI) utilized a semi-LASER sequence (TR/TE 2000/30-35 ms; flip angle 90°; FOV 160 × 160 mm; voxel size 10 × 10 × 15-20 mm). PRESS excitation was selected to encompass a grid of 8 rows } 8 columns on T2-weighted images.

### Imaging Processing

All other sequences were co-registered to the T2-weighted images in each subject. The coregistration was performed using the linear image registration tool (FLIRT) functions [18] in Oxford Centre for Functional MRI of the Brain (FMRIB) Software Library (FSL) v5.0.0 (Oxford, UK) [19]. After coregistration, DSC processing and leakage correction was performed with the NordicICE software (NordicNeuroLab, Bergen, Norway), in which the arterial input function was automatically defined. The relative cerebral blood volume (rCBV), mean transit time (MTT) and relative cerebral blood flow (rCBF) maps were calculated. DTI images were processed using the diffusion toolbox (FDT) in FSL [20], during which normalization and eddy current correction were performed. The decomposition of processed DTI images into isotropic component (p) and anisotropic component (q) was performed using the previously described method [12].

### Regions of Interest

Tumor regions of interest (ROIs) were manually delineated on the post-contrast T1 and FLAIR images using an open-source software 3D slicer v4.6.2 (https://www.slicer.org/) [21]. The delineation was independently performed by a neurosurgeon with > 8 years of experience (CL), and a researcher with > 4 years of brain tumor image analysis experience (NRB), and then reviewed by a neuroradiologist with > 8 years of experience (TM). Nonenhancing ROI, defined as the non-enhancing (NE) region outside of contrast-enhanced (CE) region, were obtained in MATLAB (MathWorks, Inc., Natick MA) by Boolean operations on contrast-enhancing and FLAIR tumor ROIs. For each individual subject, normal-appearing white matter was drawn manually in the contralateral white matter as normal controls. Each voxel value in the tumor ROI was normalized by dividing it by the mean voxel value of the contralateral normal-appearing white matter. Inter-rater reliability testing was performed using Dice similarity coefficient scores.

### Histogram Features

Histogram analysis was performed in the Statistics and Machine Learning Toolbox of MATLAB (version 2016a). All image maps, rCBV, MTT and rCBF from the perfusion images and DTI-p and DTI-q from the diffusion images, were analyzed separately. The contrast-enhancing (CE) and non-enhancing (NE) regions of interest in each map were analyzed as independent regions. Histograms were constructed using 100 bins. A total of 10 histogram features was calculated from each histogram, including mean, standard deviation, median, mode, skewness, kurtosis, and 5th, 25th, 75th, 95th percentiles of the histogram. Therefore, altogether 100 histogram features were generated from the perfusion and diffusion maps of each subject.

### Multi-view Feature Selection and Multi-view Clustering

The study design is summarized in Figure 1. Patient clustering was performed using a multiview late integration methodology called Multi-View Biological Data Analysis (MVDA), implemented in R and publicly available from GitHub (https://github.com/angy89/MVDA). Late integration methodologies allow analyzing each view independently and then merging the results [22],. For each case, we treated four categories of feature sets (CE-diffusion, NE-diffusion, CE-perfusion, NE-perfusion) as individual views to maximize the characterization of the tumor, considering the histogram features were extracted from multiple imaging modalities and regions. The analysis was divided into multiple steps: I. In order to reduce the dimensionality and remove noisy information, the features were first clustered using the hierarchical ward clustering method for each view. The number of feature clusters was determined by the previously proposed VAL index [7]. Clustering solutions with high correlation within each cluster and low correlation between the clusters were preferred. The number of features was reduced by selecting the centroids of the feature clusters which represent the features of each view. II. For each view, the patients were clustered by applying a hierarchical ward clustering method using the features selected from previous step. The number of patient clusters was also determined by the VAL index. III. The clustering results of each view were integrated in a late integration method. The vector of clustering assignment of each view was transformed in a binary membership matrix, with the patients on the rows and the clustering on the columns. These matrices were transposed and stacked vertically to create a bigger matrix X with L rows (the clusters) and N columns (the patients). This matrix was then factorized in order to obtain two matrices P (with L rows and k columns) and H (with k rows and N columns). The difference between X and PH was as minimum as possible. In this settings H represented the membership matrices of the N patients to the final multi-view clusters. The number of multi-view cluster was set to 2 to dichotomize patients into subgroups with better and worse survivals.

**Figure 1.**
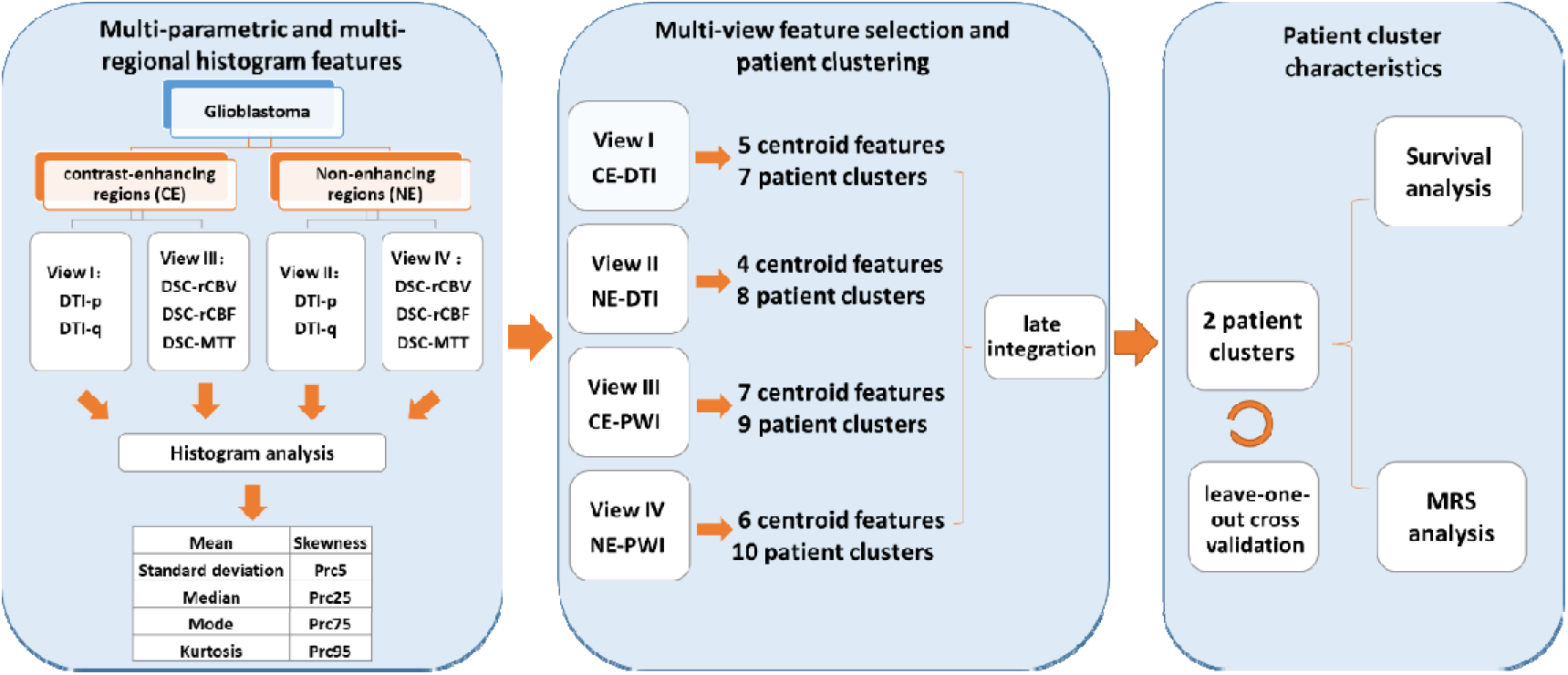
Study design. DTI-p and DTI-q maps were generated from diffusion tensor imaging (DTI). The relative cerebral blood volume (rCBV), mean transit time (MTT) and relative cerebral blood flow (rCBF) maps were generated from dynamic susceptibility contrast (DSC) imaging. Histogram features were extracted from the multiple modalities and regions (contrast-enhancing and non-enhancing), then were separated into four independent views. Each view was first clustered to select the centroid features, which later were used to cluster patients. The resulting clusters from each view were integrated to generate two patient clusters. The clusters were assessed using leave-one-out cross validation and survival analysis. The metabolic signatures of the two clusters were compared.

### Leave-one-out Cross Validation of the Clustering

A leave-one-out cross-validation (LOOCV) procedure was applied for constructing and validating the patient clusters. All steps of MVDA approach was repeated by leaving one patient out of the cohort at each repetition. The consensus analysis was performed in the 80 clustering results obtained from the LOOCV approach. An 80 × 80 co-occurrence consensus clustering matrix M was created, where M (*i, j*) indicating percentage of times that the patients *i* and *j* were clustered together across the 80 dataset perturbations.

### Feature Ranking

The importance of each selected feature was calculated using receiver operator characteristics (ROC) curve analysis, on the final multi-view clustering result. The analysis was performed by using an R package ‘Caret’[23]. The final patient clustering results was used to train a learning vector quantization (LVQ) model and calculates a resampling based performance measure. The importance of each feature was ranked according to the area under the ROC curves (AUC).

### Multivoxel MRS Processing

MRS data were processed using LCModel (Provencher, Oakville, Ontario). All the concentrations of metabolite were calculated as a ratio to creatine (Cr). All relevant spectra from CSI voxels of interest were assessed for artefacts using previous criteria [24]. The values of the Cramer–Rao lower bounds were used to evaluate the quality and reliability of CSI data and values with standard deviation (SD) > 20% were discarded. To account for the difference in spatial resolution, the T2-space pixels were projected to CSI space according to their coordinates using MATLAB. The proportion of T2-space tumor pixels occupying each CSI voxel was calculated. A selection rule was applied that only those CSI voxels were included when the proportion of tumor pixels were over 50%. Only CSI voxels contacting more than 50% tumor T2-voxels were included for further analysis. The weight of each CSI voxel was taken as the proportion of the tumor pixels in that CSI voxel. The summed weighted value was used as final metabolic value of the tumor ROI.

### Statistical Analysis

All statistical analyses were performed in RStudio v3.2.3. Non-normal distributed CSI data were compared with Wilcoxon rank sum test using Benjamini-Hochberg procedure for controlling the false discovery rate in multiple comparisons. Kaplan-Meier and Cox proportional hazards regression analyses were performed to evaluate patient survival. For Cox proportional hazards regression, all the confounders, including IDH-1 mutation status, MGMT methylation status, sex, age, extent of resection and contrast-enhancing tumor volume were considered. For the Kaplan-Meier analysis, each feature was dichotomized for OS and PFS before the log-rank test by using optimal cutoff values calculated by ‘surv_cutpoint’ function in the R Package “survminer” Patients who were alive at the last known follow-up were censored. Significance was accepted at a two-sided significance level of alpha < 0.05.

## Results

### Patients and regions of interest

The mean age of the patients was 57.4 years (range 22-73 years). The median overall survival of the patients was 461 days (range 52-1259 days) and the median progression-free survival was 264 days (range 37-1130 days). Patient characteristics were summarized in Table 1. Inter-rater reliability testing of regions of interest (ROIs) showed excellent agreement between the two raters, with Dice scores 0.85 ± 0.10 (mean ± standard deviation [SD]) for contrast-enhancing and 0.86 ± 0.10 of FLAIR ROIs respectively. The volumes of contrast-enhancing and non-enhancing ROIs were 49.7 cm^3^ ± 28.1cm^3^ and 64.7 cm^3^ ± 48.3 cm^3^ respectively.

**Table 1.** Clinical characteristics

| **Table 1. Clinical characteristics** |  |
| --- | --- |
| Variable | Patient Number |
| **Age at diagnosis** |  |
| <60 | 35 |
| ≥60 | 45 |
| **Sex** |  |
| Male | 58 |
| Female | 22 |
| **Extent of resection (of enhancing tumor)** |  |
| Complete resection | 56 |
| Partial resection | 22 |
| Biopsy | 2 |
| **MGMT-methylation status\*** |  |
| Methylated | 23 |
| Unmethylated | 31 |
| **IDH-1 mutation status** |  |
| Mutant | 7 |
| Wild-type | 73 |
| **Tumor volumes(cm<sup>3</sup>)<sup>#</sup>** |  |
| Contrast-enhancing | 49.7 ± 28.1 |
| Non-enhancing | 64.7 ± 48.3 |
| **Survival (days)** |  |
| Median OS (range) | 461 (52-1259) |
| Median PFS (range) | 264(37-1130) |
| *MGMT-methylation status unavailable for 26 patients; <sup>#</sup>mean ± SD of original data. SD: standard deviation; MGMT: O-6-methylguanine-DNA methyltransferase; IDH-1: Isocitrate dehydrogenase 1; cm: centimeters; OS: overall survival; PFS: progression-free survival. |  |

**Table 2.** Histogram features

| Table 2. Histogram features |  |
| --- | --- |
| Feature | Description |
| Mean | mean intensity of voxels |
| Standard deviation (SD) | standard deviation of intensity of voxels |
| Median | median intensity of voxels |
| Mode | most common intensity of voxels |
| Skewness | measure of the symmetry of the histogram |
| Kurtosis | measure of the tail of the histogram |
| 5th percentile (Prc5) | fifth percentile of the intensity histogram |
| 25th percentile (Prc25) | twenty-fifth percentile of the intensity histogram |
| 75th percentile (Prc75) | seventy-fifth percentile of the intensity histogram |
| 95th percentile (Prc95) | ninety-fifth percentile of the intensity histogram |

### Feature Selection and Multi-view Clustering

For the four different views, 5, 4, 7 and 6 centroid features were respectively selected by the algorithm and listed in Table 3 and demonstrated by Figure2. Using these centroid features and the optimal number of clusters, patients were firstly divided into 7, 8, 9 and 10 clusters respectively in the four views using hierarchical ward clustering. Late integration of the four views yielded a final clustering of two patient subgroups, with 53 and 27 patients in each subgroup respectively. Kaplan-Meier analysis showed significance for overall survival (Log-rank, p = 0.020) and progression-free survival (Log-rank, p < 0.001) (Figure 5A &5B). Significant difference was also found in the overall survival (p = 0.007, HR = 0.32) and progression-free survival (p < 0.001, HR = 0.33) of the two patient subgroups, using multivariate Cox proportional hazards regression,

**Figure 2.**
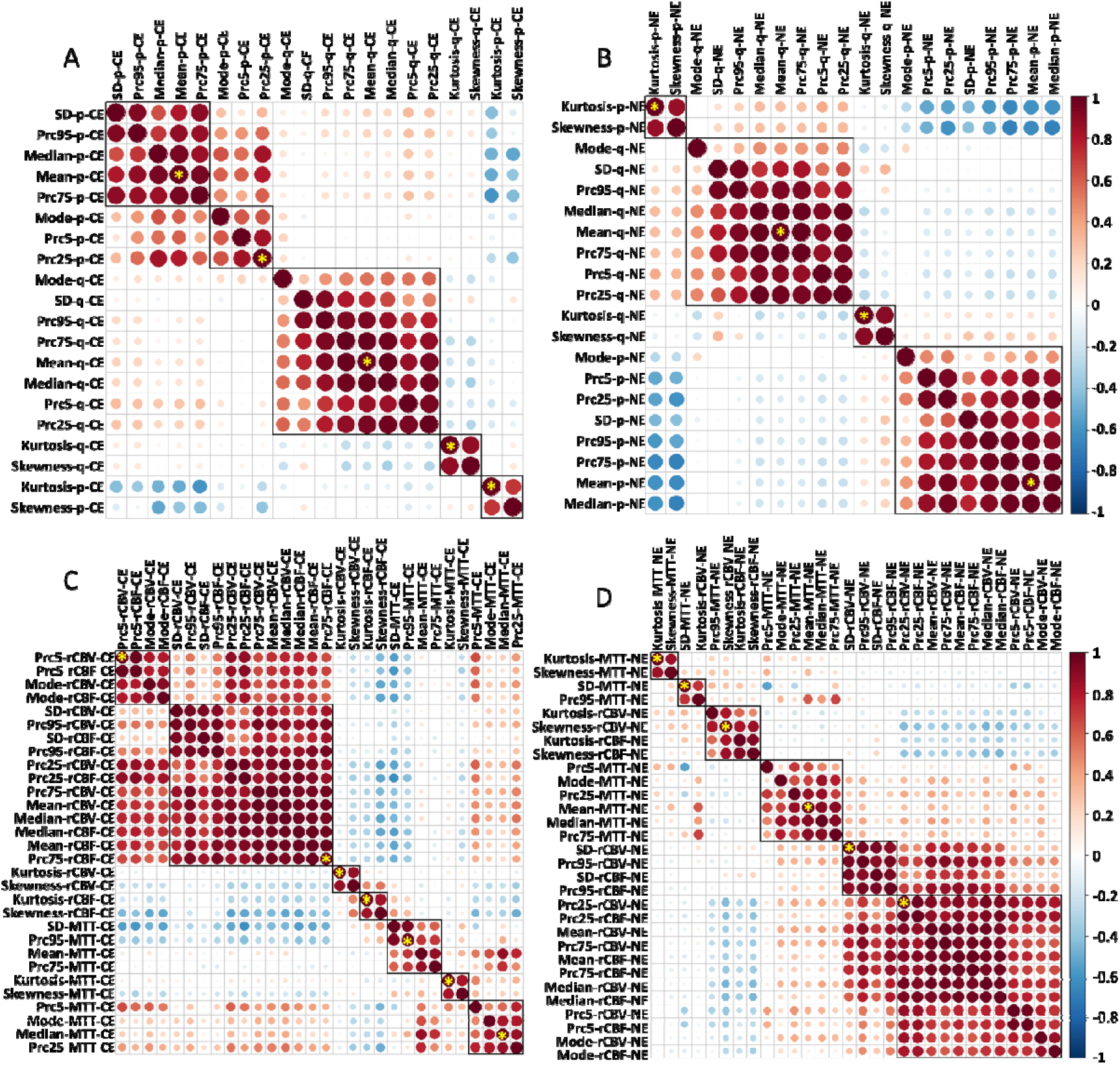
Feature clustering and centroid features. In each view, all features were clustered using the hierarchical ward clustering method. The centroid features (marked by yellow stars) were selected to represent each view. A: view 1 (perfusion histogram features in the contrast-enhancing regions); B: view 2 (perfusion histogram features in the non-enhancing regions); C: view 3 (diffusion histogram features in the contrast-enhancing regions); D: view 4 (diffusion histogram features in the non-enhancing regions).

**Figure 5.**
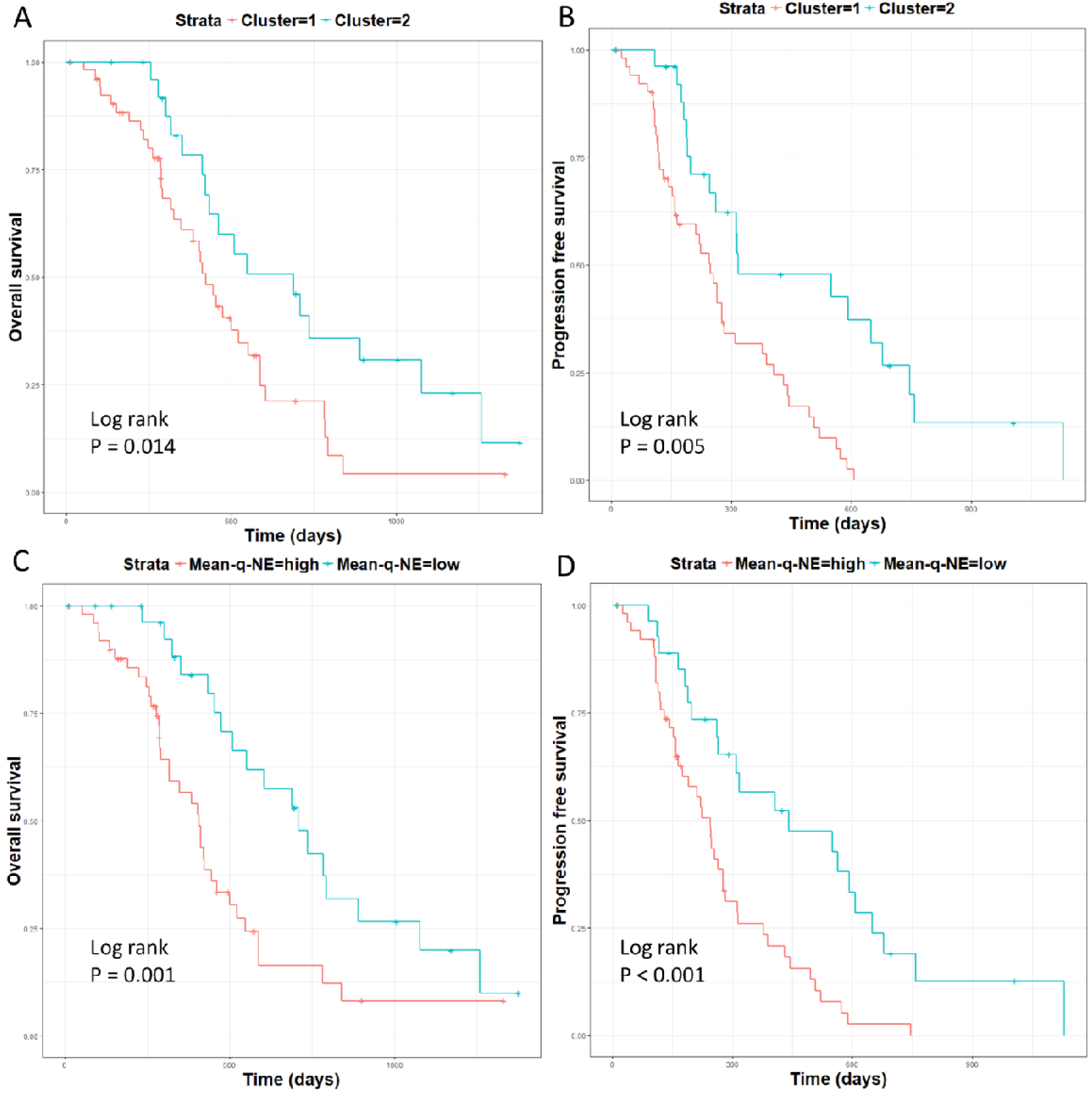
Kaplan-Meier plots of survival analysis. Log-rank test showed patient cluster 2 displayed better OS (p = 0.020) (A) and PFS (p < 0.001) (B). Higher man value of DTI-q in the non-enhancing region (Mean-q-NE) was associated with a worse OS (p = 0.002) (C) and PFS (p < 0.001) (D).

**Table 3.**
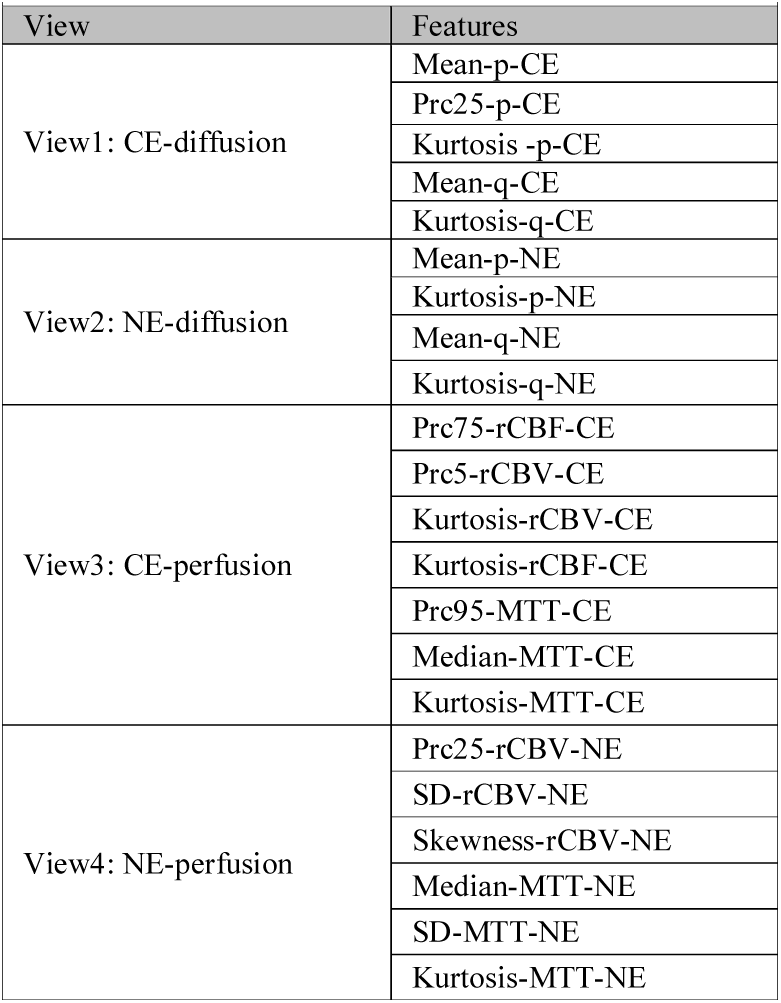
Features selected from multi-view clustering

After the leave-one-out cross validation, the co-occurrence consensus clustering matrix was computed and showed that the two patient clusters generated from the unsupervised clustering were stable. The mean value of the co-occurrence consensus clustering matrix were 0.79 for patient cluster 1 and 0.68 for patient cluster 2. The co-occurrence consensus clustering results are demonstrated in Figure 3.

**Figure 3.**
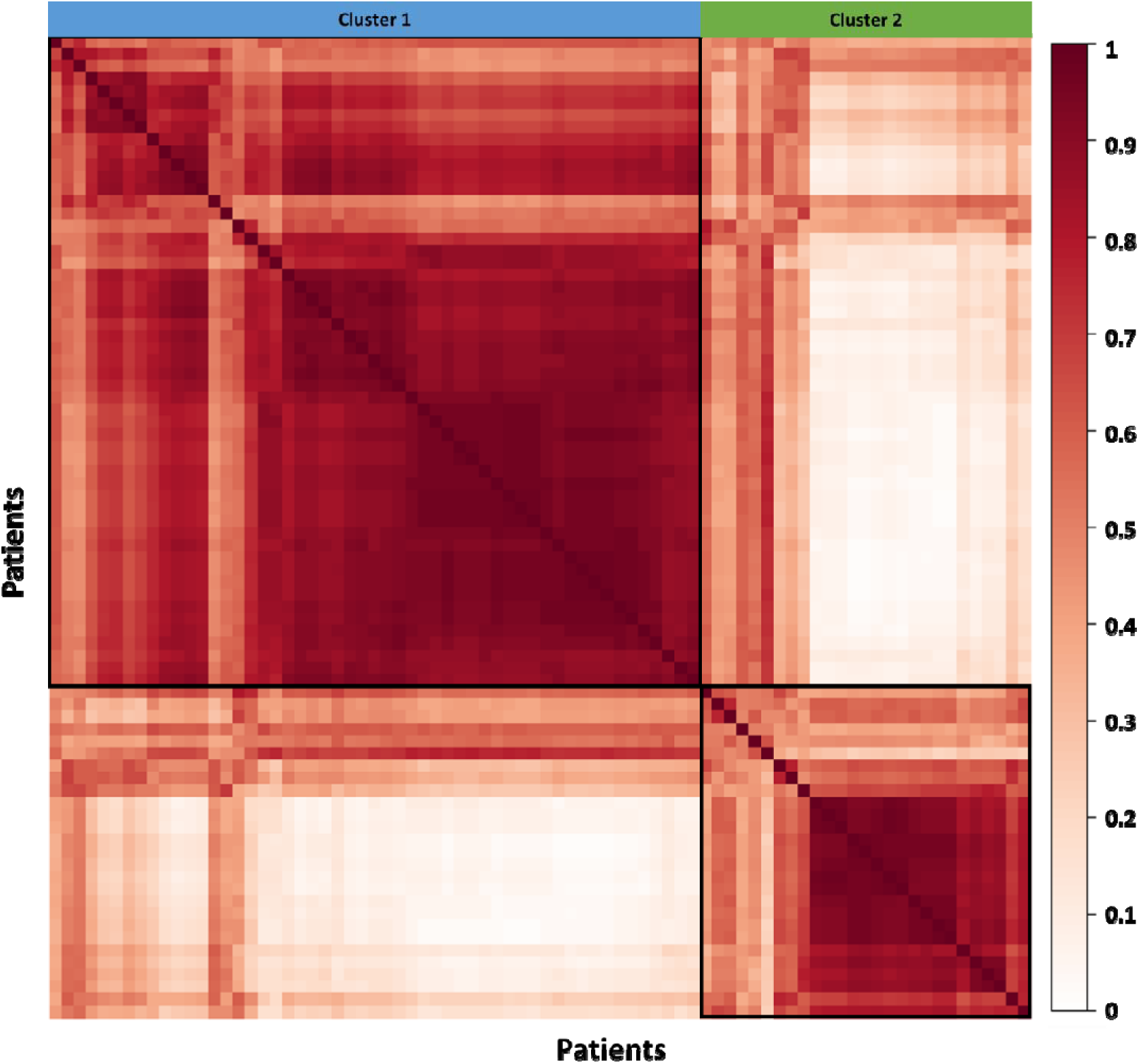
Patient clustering and co-occurrence consensus clustering matrix. After multi-view clustering, consensus analysis was performed based on the 80 clustering results obtained after the leave-one-out cross validation. The mean value of the co-occurrence consensus clustering matrix was 0.79 for patient cluster 1 and 0.68 for patient cluster 2.

### Feature ranking

To assess the importance of features, they were ranked according to the AUC, listed in Table 6 and demonstrated by Figure 4. The top 5 most important features were: Mean-p-NE (mean value of DTI-p in the non-enhancing tumor regions) (AUC = 0.867), Mean-q-NE (mean value of DTI-q in the non-enhancing tumor regions) (AUC = 0.804), Prc25-rCBV-NE (25th percentile of rCBV in the non-enhancing tumor regions) (AUC = 0.776), Kurtosis-p-NE (kurtosis of DTI-p histogram in the non-enhancing tumor regions) (AUC = 0.748), Mean-q-CE (mean value of DTI-q in the contrast-enhancing tumor regions) (AUC = 0.738).

**Figure 4.**
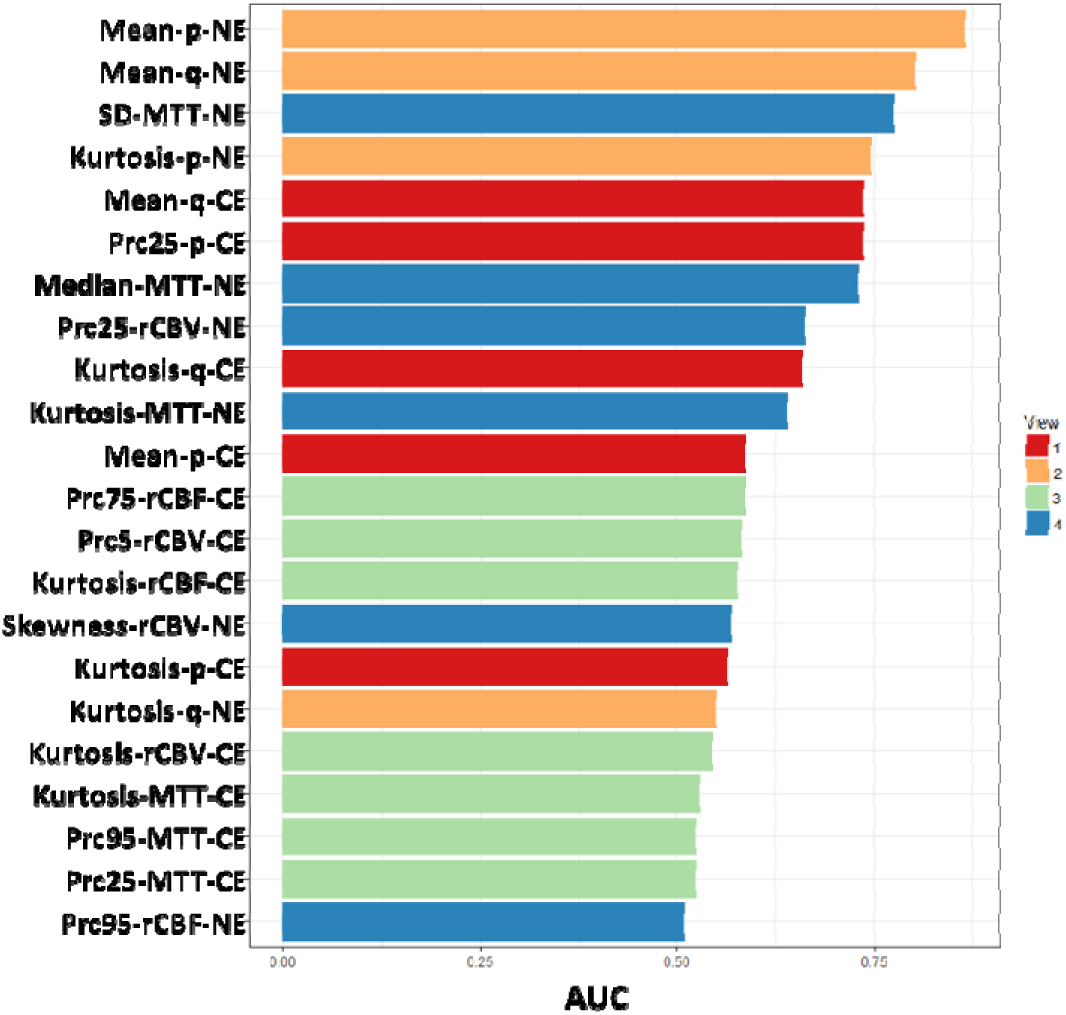
Feature ranking. The selected features were ranked according to the area under the receiver operator characteristics curves (AUC) to assess their importance.

### Multivariate survival analysis of individual features

The multivariate survival modeling of PFS and OS were tested in 78 patients for whom all confounders including IDH-1 mutation and MGMT methylation status were available. The results of the multivariate Cox-regression analysis are shown in Table 4 and demonstrated by Figure 5C & 5D. Four diffusion histogram features significantly contributed to patient survivals. Specifically, higher Mean-q-NE contributed to a worse OS (HR = 1.40, CI: 1.05 0. 86, p = 0.020) and worse PFS (HR = 1.36, CI: 1.03-1.79, p = 0.031).

**Table 4.**
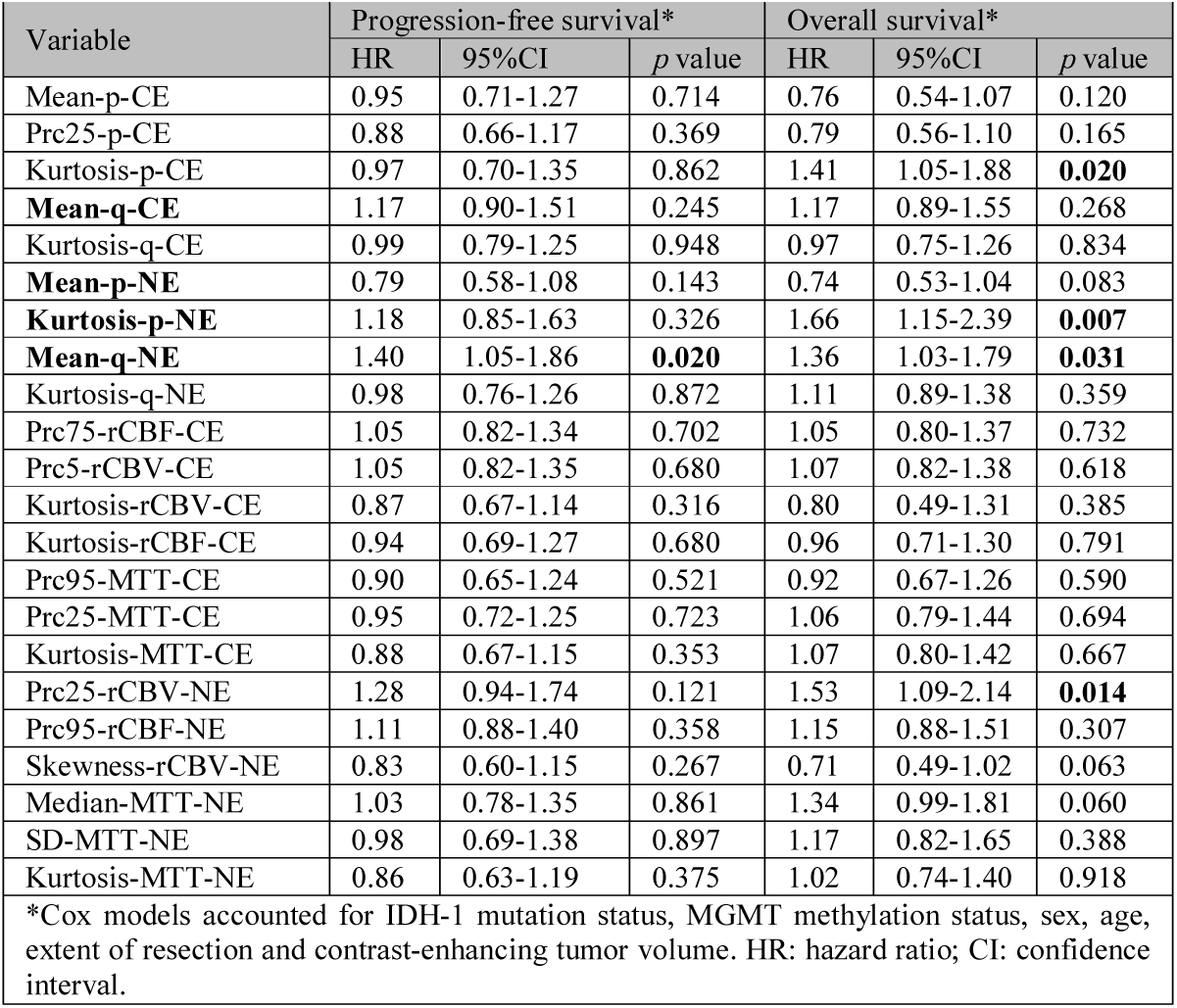
Cox multivariate modeling of survivals

**Table 5.** Metabolic statistics

| Metabolite | Contrast-enhancing tumor region |  |  |  |  | Non-enhancing tumor region |  |  |  |  |
| --- | --- | --- | --- | --- | --- | --- | --- | --- | --- | --- |
|  | Cluster 1 |  | Cluster 2 |  | p | Cluster 1 |  | Cluster 2 |  | p |
| | Mean ± SD | 95% CI | Mean ± SD | 95% CI | | Mean ± SD | 95% CI | Mean ± SD | 95% CI | |
| Cho/NAA | 0.60 ± 0.32 | 0.50-0.69 | 0.49 ± 0.26 | 0.38-0.60 | 0.091 | 0.54 ± 0.50 | 0.38-0.70 | 0.45 ± 0.19 | 0.37-0.53 | 0.689 |
| Cho/Cr | 0.70 ± 0.24 | 0.63-0.77 | 0.63 ± 0.17 | 0.55-0.70 | 0.244 | 0.44 ± 0.13 | 0.40-0.48 | 0.40 ± 0.14 | 0.34-0.46 | 0.149 |
| NAA/Cr | 0.76 ± 0.35 | 0.66-0.86 | 0.46 ± 0.27 | 0.35-0.58 | **<0.001** | 0.97 ± 0.32 | 0.86-1.07 | 0.76 ± 0.30 | 0.64-0.89 | **0.013** |
| GSH/Cr | 0.31 ± 0.28 | 0.23-0.40 | 0.40 ± 0.36 | 0.23-0.57 | 0.245 | 0.34 ± 0.19 | 0.27-0.41 | 0.39 ± 0.27 | 0.26-0.51 | 0.687 |
| Glu/Cr | 1.57 ± 0.86 | 1.31-1.83 | 1.54 ± 1.08 | 0.95-1.61 | 0.648 | 1.04 ± 0.41 | 0.89-1.18 | 1.37 ± 0.60 | 1.11-1.62 | **0.037** |
| Glx/Cr | 1.89 ± 1.11 | 1.56-2.23 | 1.85 ± 1.29 | 1.31-2.40 | 0.651 | 1.41 ± 0.58 | 1.20-1.62 | 1.98 ± 1.04 | 1.54-2.42 | **0.027** |
| mIn/Cr | 1.37 ± 0.88 | 1.12-1.62 | 1.28 ± 0.77 | 1.08-1.99 | 0.657 | 1.14 ± 0.41 | 1.00-1.27 | 1.16 ± 0.38 | 1.00-1.32 | 0.924 |
| Lac/Cr | 6.37 ± 5.73 | 4.67-8.07 | 4.81 ± 4.14 | 2.93-6.69 | 0.448 | 1.13 ± 0.98 | 0.75-1.52 | 1.45 ± 2.62 | 0.11-2.80 | 0.087 |

Choline; NAA: N-acetyl aspartate; GSH: glutathione; Glu: glutamate; Glx: glutamate + glutamine; mIn: myo-inositol; Lac: lactate; CI: confidence interval.

### Multivoxel MR Spectroscopy

Due to the abovementioned MRS analytic rules excluding CSI voxels containing less than 50% tumor, CSI data were missing in four patients. Our results showed N-acetylaspartate/creatine (NAA/Cr) ratio of the cluster 1 was significantly higher than in cluster 2, both in the CE region (p < 0.001) and NE region (p = 0.013). In the NE region, both glutamate/Cr (Glu/Cr) ratio and glutamate + glutamine/Cr (Glx/Cr) of the cluster 1 was significantly lower than cluster 2 (p = 0.037, and 0.027 respectively). No other metabolites showed significant differences. The metabolic results of the two patient clusters are demonstrated by Figure 6.

**Figure 6.**
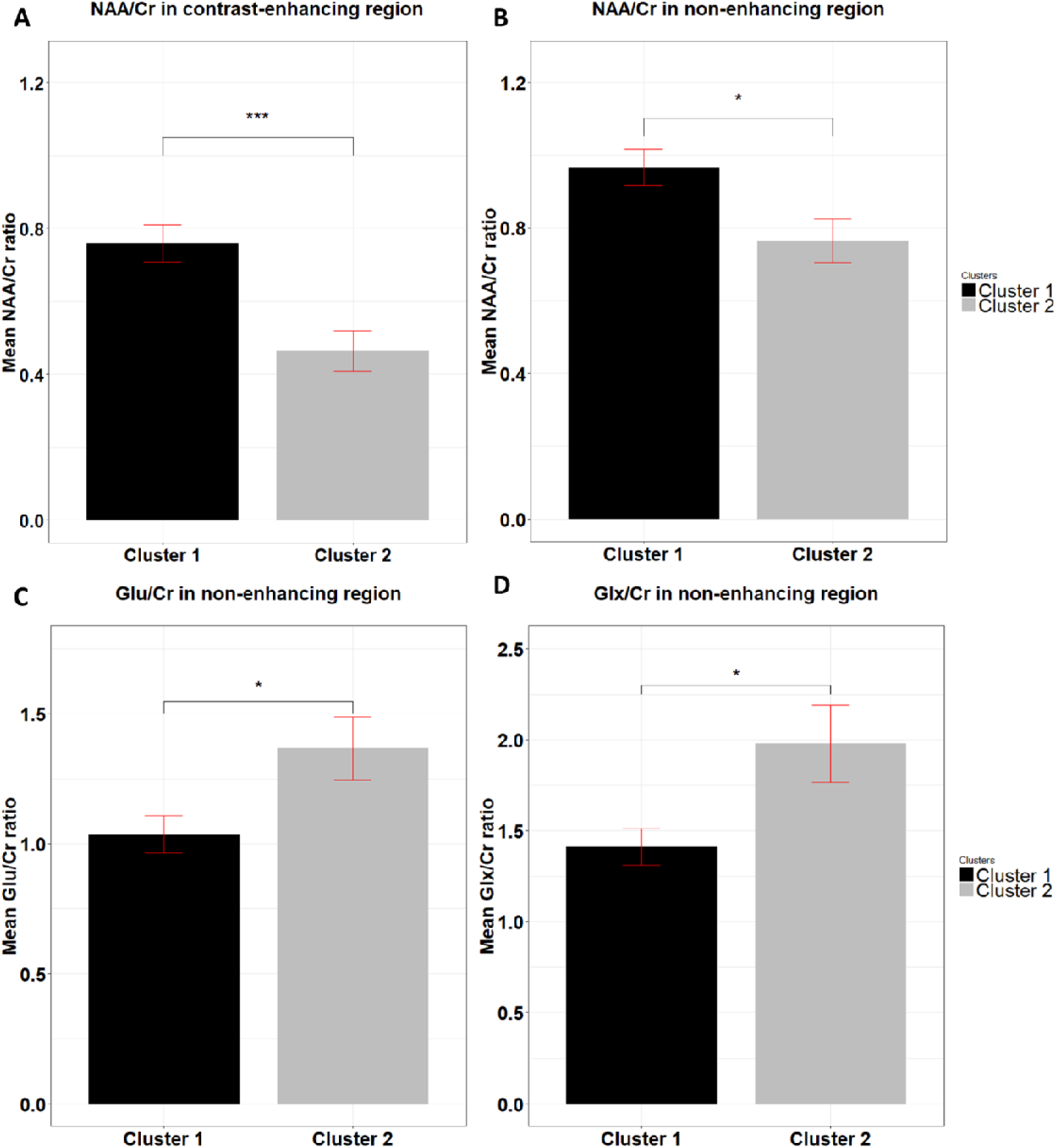
MRS results. N-acetylaspartate/creatine (NAA/Cr) ratio in cluster 1 was significantly higher than in cluster 2, both in the contrast-enhancing (CE) region (p < 0.001) (A) and non-enhancing (NE) region (p = 0.013) (B). In the NE region, cluster 1 showed significantly higher glutamate/Cr (Glu/Cr) ratio (p = 0.037) (C) and glutamate + glutamine/Cr (Glx/Cr) ratio (p = 0.027) (D) than cluster 2.

## Discussion

This study demonstrated that integrating multi-parametric and multi-regional MRI histogram features may help to stratify patients; the histogram features extracted from diffusion tensor imaging are particularly useful for predicting patient outcomes, as demonstrated by the multivariable Cox regression analysis.

Histogram analysis of quantitative MRI offers a volumetric characterization of the tumor heterogeneity [25, 26]. Several studies have investigated the utility of MRI histogram features in tumor heterogeneity evaluation and survival prediction [27, 28]. However, only limited studies have included both perfusion and diffusion tensor imaging variables. In one of these studies, diffusion and perfusion imaging features only showed marginal prognostic values [27]. The multi-view approach we used can offer the advantage of the parallelized selection of features from different modalities. Being a late integration approach, the analyses on each view are independent and can be analyzed in parallel. It can also avoid the representation issues, since the clustering results generate from each independent view are the inputs to the final integration algorithms [7]. With this algorithm, we finally separated patients into two groups, which showed survival difference. Our results may suggest that integrate these imaging features properly is crucial in patient stratification.

We ranked the histogram features according to their importance in the patient clustering and tested them in the multivariate Cox regression model. Consistent with previous studies [29], our results showed that DTI histogram parameters were useful in survival prediction. Moreover, our results also suggested that diffusion tensor imaging have higher prognostic values over perfusion imaging. In the previous results, rCBV was reported to have predicative values for patient survivals [30]. Interestingly, although our results showed rCBV was useful in patient clustering, it failed to show a significant impact on survival in the multivariate analysis, which may challenge its robustness.

Glioblastoma is recognized to preferentially migrate along the white matter tracts which may provide a ‘fast track’ for tumor infiltration and lead to the increased anisotropic movement. The diffusion of water molecules in the tumor core and peritumoral brain tissue is consequently altered. Our results showed the mean value of DTI-q may contribute to a worse patient survival. Further, as NAA is a marker of neurons, the increased NAA level in the worse patient group may suggest the more infiltrated tumor phenotype, which corresponds to the increased anisotropic movement of water molecules revealed by the elevated DTI-q value. Glutamate is a key neurotransmitter in the brain and is central for the neuronal functions [31]. Previous MR spectroscopy study showed increased level of glutamate in oligodendroglioma [32], and in the necrotic regions of glioblastoma [33]. These studies suggested that the high extracellular levels of glutamate may be caused by the structural destruction in the tumor [31], which corresponds to our current results, showing that cluster 2 may have more destructed neuronal fibers (lower NAA/Cr and higher Glx/Cr levels).

Our study has some limitations. Firstly, although we used the leave-one-out cross validation, the patient population reported is from a single center. Secondly, although previous studies have validated the histological correlates of DTI-p and DTI-q by image-guided biopsies, our current findings need further biological validation. Lastly, as the ^1^H-MRS voxels were larger than T2 space voxels, we had fewer patients with lactate data available and the multivariate analysis was done with a smaller sample size.

In conclusion, our results showed that the multi-view clustering method can provide an effective approach of integrating multiple quantitative MRI features. The histogram features selected from the proposed approach may be used as potential imaging markers in personalized treatment strategy and response determination.

